# The Human Thalamus is an Integrative Hub for Functional Brain Networks

**DOI:** 10.1101/056630

**Authors:** Kai Hwang, Maxwell Bertolero, William Liu, Mark D’Esposito

## Abstract

The thalamus is globally connected with distributed cortical regions, yet the functional significance of this extensive thalamocortical connectivity remains largely unknown. By performing graph-theoretic analyses on thalamocortical functional connectivity data collected from human participants, we found that the human thalamus displays network properties capable of integrating multimodal information across diverse cortical functional networks. From a meta-analysis of a large dataset of functional brain imaging experiments, we further found that the thalamus is involved in multiple cognitive functions. Finally, we found that focal thalamic lesions in humans have widespread distal effects, disrupting the modular organization of cortical functional networks. This converging evidence suggests that the human thalamus is a critical hub region that could integrate heteromodal information and maintain the modular structure of cortical functional networks.

## Introduction

The cerebral cortex has extensive anatomical connections with the thalamus. Every cortical region receives projections from the thalamus, and in turn sends outputs to one or multiple thalamic nuclei (Jones, 2001). The thalamus is one of the most globally connected neural structures (Cole et al., 2010; Crossley et al., 2013); thalamocortical projections relay nearly all incoming information to the cortex, as well as mediate cortico-cortical communication (Sherman and Guillery, 2013). The mammalian brain can therefore be conceptualized as a thalamocortical system. Thus, full insight into brain function requires knowledge of the organization and properties of thalamocortical interactions.

The thalamus can be divided into two types of nuclei: first order and higher order thalamic nuclei (Sherman, 2007). First order thalamic nuclei, such as the lateral geniculate nucleus (LGN), ventral posterior (VP) and ventral lateral (VL) nuclei, receive inputs from ascending sensory pathways or other subcortical brain regions. In contrast, higher-order thalamic nuclei, such as the mediodorsal (MD) and the pulvinar nuclei, receive inputs predominately from the cortex. More than half of the thalamus comprises higher order thalamic nuclei, which have both reciprocal and non-reciprocal connections with multiple cortical regions. These connectivity profiles suggest that in addition to relaying sensory and subcortical information to the cortex, another principle function of the thalamus is to mediate the transfer of information between cortical regions through cortico-thalamo-cortical pathways (Sherman, 2016).

How a brain region processes and communicates information can be inferred by its connectivity pattern (Passingham et al., 2002). Graph-theoretic network analysis of resting-state functional MRI (rs-fMRI) data is well suited for exploring the network properties of the thalamocortical system (Bullmore and Sporns, 2009). Functional connectivity analyses of rs-fMRI data measures correlations of spontaneous fluctuations in blood oxygenation level dependent (BOLD) signals, which are not a direct proxy for anatomical connectivity but are largely constrained by anatomical connections (Hermundstad et al., 2013; Honey et al., 2009). Functional connectivity between two brain regions likely represents the phase-locking of the two regions' low-frequency oscillations or coherent activity of high-frequency neuronal activity (Scholvinck et al., 2010; Wang et al., 2012).

Previous functional connectivity analyses of rs-MRI data have consistently revealed a modular organization of the human cerebral cortex, indicating that the cortex is composed of several specialized functional networks (Power et al., 2011; Yeo et al., 2011). Each of these networks is potentially involved in executing a discrete set of cognitive functions (Bertolero et al., 2015; Smith et al., 2009). Graph-theoretic measures can be used to quantify topographic properties of each brain region and make inferences on each region's network functions (Sporns et al., 2007). For instance, a brain region with many within-network connections has a strong “provincial hub” property (Guimera and Amaral, 2005), presumably to promote within-network interactions for executing specialized functions of the network; whereas a brain region with many between-network connections has a strong “connector hub” property, presumably to mediate interactions between functional networks. Connector and provincial hubs have distinct contributions to modular organization. For example, a lesion study showed that damage to connector hubs, but not provincial hubs, causes more severe disruption of the network's modular organization (Gratton et al., 2012), suggesting that focal lesions to connector hubs can have a widespread impact on network organization when between-network connections are disrupted. Finally, cortical connector hubs are involved in multiple cognitive tasks (Cole et al., 2013; Warren et al., 2014; Yeo et al., 2015), and exhibit increased activity when multiple functional networks are engaged by a behavioral task (Bertolero et al., 2015). These findings suggest that connector hubs are capable of multimodal and integrative processing through their extensive between-network connectivity (van den Heuvel and Sporns, 2013).

The thalamus has been largely ignored in studies of brain network organization, and the topographic properties of the thalamocortical system are largely unknown. Previous graph-theoretic studies of functional brain networks often exclude subcortical structures, or examine the thalamus with gross or no subdivisions. However, given its complex structure with multiple distinct nuclei, the thalamus is likely not uniformly interacting with the cortex. Different thalamic subdivisions have distinct structural connectivity with the cortex, and thus functional connectivity with the cortex (Arcaro et al., 2015; Behrens et al., 2003; Yuan et al., 2015). Traditionally, it is proposed that each thalamic subdivision functionally connects with cortical regions that belong to the same functional network for partially closed-loop, modality-selective processes (Alexander et al., 1986). Based on this hypothesis, thalamic subdivisions should exhibit strong provincial hub (within-network) properties (Figure 1A). Alternatively, if a thalamic subdivision functionally interacts with cortical regions from multiple functional networks, it should exhibit strong connector hub (between-network) properties (Figure 1B). These hypotheses are not mutually exclusive—the thalamus could contain subdivisions that are involved in both modality-selective and multimodal, integrative processes.

**Figure 1.**
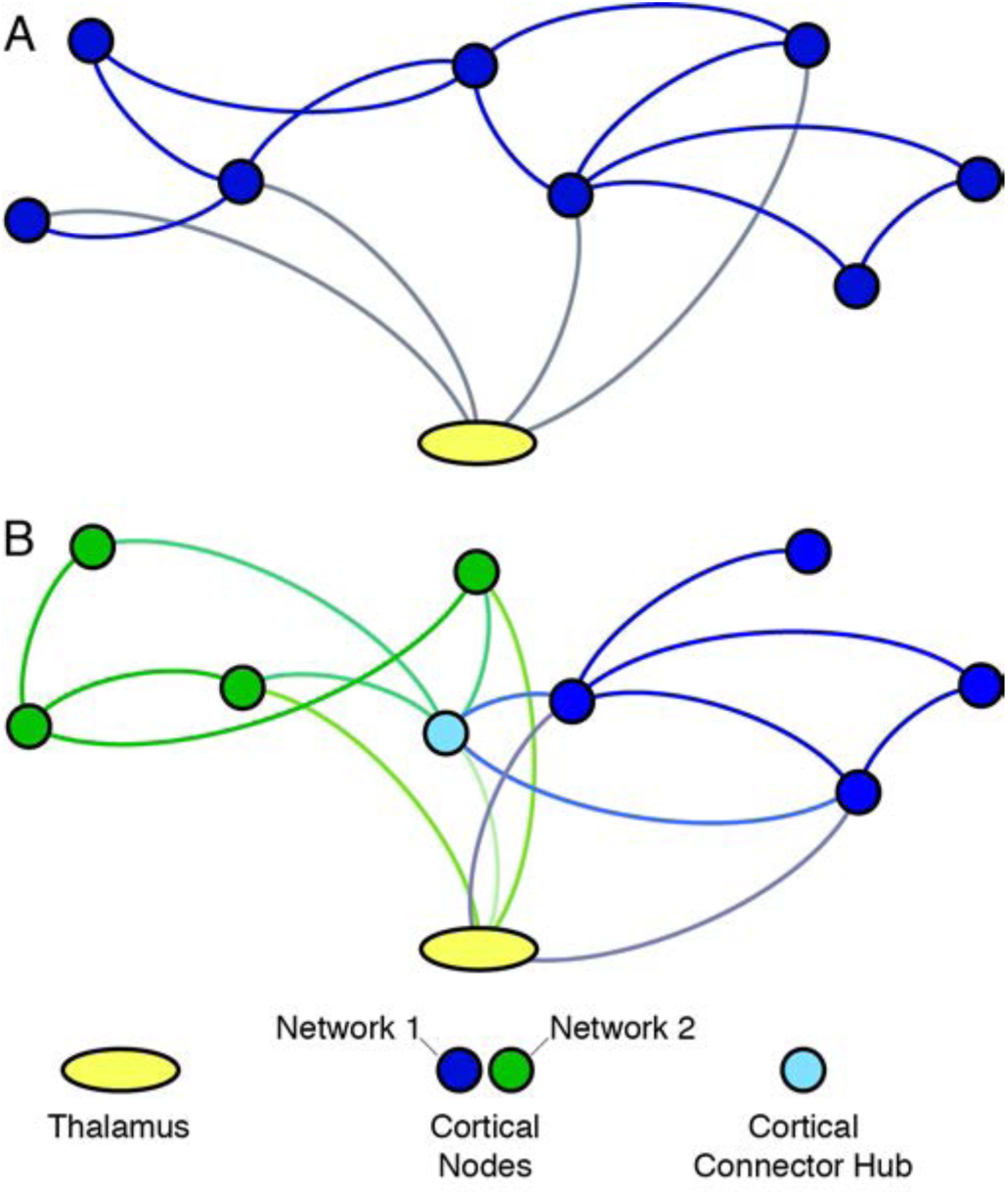
Thalamus mediating cortico-cortical communication for functional brain networks. (A) As a provincial hub, the thalamus is connected with cortical regions that belong to the same cortical functional network (represented in solid blue circles). (B) As a connector hub, the thalamus is connected with cortical regions in multiple cortical functional networks (one network colored in blue and the other in green).

The goal of this study was to elucidate the thalamus's network topological role in functional brain networks. To measure network properties of thalamocortical functional connectivity, we performed graph theoretic network analyses on rs-fMRI data collected from healthy human participants. To relate network topology to cognitive functions, we analyzed task-related activity of the thalamus using a meta-analysis of 10,449 functional neuroimaging experiments from the BrainMap database (Laird et al., 2005; Yeo et al., 2015). Finally, we examined the thalamus's contribution to cortical network organization by analyzing rs-fMRI data from human patients with focal thalamic lesions.

## Results

### Identification of Cortical Networks

To identify cortical functional networks, we first measured functional connectivity matrices between 333 cortical ROIs (Gordon et al., 2016), then performed a network partition analysis to estimate cortical network organization (see Methods). We found that the cerebral cortex can be decomposed into 9 functional networks (Figure 2A), an organization scheme largely similar to previous studies (Bertolero et al., 2015; Gordon et al., 2016; Power et al., 2011; Yeo et al., 2011).

**Figure 2.**
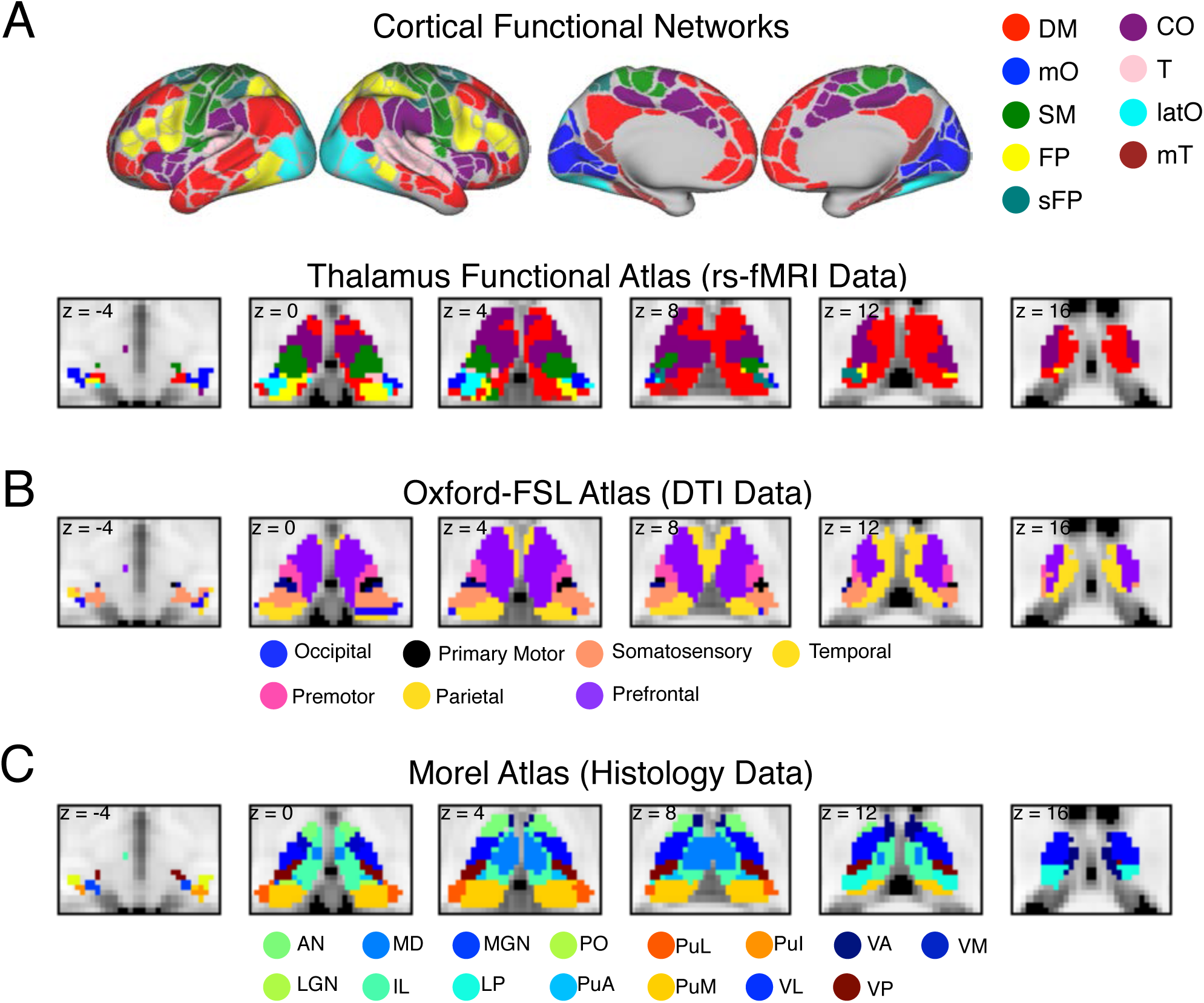
Cortical functional networks and thalamic atlases. (A) Cortical functional networks and thalamic parcellation derived from functional connectivity analyses between the thalamus and each cortical network using rs-fMRI data. Network abbreviations (based on its most predominant anatomical location): default mode (DM), medial occipital (mO), somato-motor (SM), fronto-parietal (FP), superior fronto-parietal (sFP), cingulo-opercular (CO), temporal (T), lateral occipital (latO), medial temporal (mT). A detailed list of the specific ROIs for each network are provided in Supplementary Table 1 (B) Structural connectivity based segmentation of the thalamus using the Oxford-FSL atlas. Each thalamic subdivision was labeled based on the cortical region it is most structurally connected with. (C) Histology based thalamic parcellation using the Morel atlas. Abbreviations for thalamic nuclei: anterior nucleus (AN), intralaminar (IL), lateral posterior (LP), lateral geniculate nucleus (LGN), medial geniculate nucleus (MGN), medial dorsal (MD), medial pulvinar (PuM), inferior pulvinar (PuI), lateral pulvinar (PuL), anterior pulvinar (PuA), posterior (Po) nuclei, ventral posterior (VP), ventral anterior (VA), ventral medial (VM), Ventral lateral (VL).

#### Parcellation of the Thalamus

Given that the thalamus can be subdivided using different approaches, we performed our analyses using three different atlases based on data from rs-fMRI, diffusion tensor imaging (DTI), and postmortem histology (Figure 2 A-C; see Methods for details). Using RS-fcMRI data, we identified thalamic subdivisions that demonstrated the strongest functional connectivity with the different cortical functional networks reported above (Figure 2A; henceforth referred to as the functional parcellation atlas). We further replicated these results with an independent dataset, and found high correspondence between datasets (normalized mutual information = 0.69, z-scored Rand coefficient =144.13, *p* < 10^−5^). The Oxford-FSL thalamocortical structural connectivity atlas (Figure 2B) subdivides the thalamus based on structural connectivity (estimated using probabilistic diffusion tractography on DTI data) to seven large cortical areas: primary motor, primary somatosensory, occipital, premotor, prefrontal (including medial and orbitofrontal cortices), parietal, and temporal cortices (Behrens et al., 2003). The Morel atlas (Figure 2C) subdivides the thalamus into smaller nuclei based on cyto- and myelo-architecture information from five postmortem brains (Krauth et al., 2010; Morel et al., 1997). We further classified each thalamic nucleus from the Morel atlas into first order or higher order thalamic nuclei (Sherman, 2016; Sherman and Guillery, 2013).

### Network Properties of Thalamocortical Functional Connectivity

To determine each thalamic subdivision's network property, we measured functional connectivity between each thalamic voxel and every cortical ROI (see Methods) to generate a thalamic voxel to cortical ROI thalamocortical network graph. Graph metrics were calculated for every thalamic voxel, and pooled across voxels for different categories of thalamic subdivisions (i.e., first order and higher order nuclei, or subdivisions within the three different thalamic atlases). We also calculated graph metrics at the level of thalamic subdivision by first averaging BOLD signals within each subdivision, and then estimated each subdivision's thalamocortical functional connectivity with cortical ROIs. Our goal was to examine the thalamus's network topological properties in the context of large-scale functional brain networks. For example, if a thalamic subdivision is found to have strong connector hub properties, how does it compare to cortical ROIs that are connector hubs? Therefore, we further calculated graph metrics for each cortical ROI by analyzing patterns of cortico-cortical functional connectivity.

#### Provincial Hub Property Analyses

Provincial hub properties can be measured by the graph theory metric called within module degree (WMD), which measures the number of within-network connections of a region, z-scored by the mean and standard deviation of within-network connections of all regions in that network (Guimera and Amaral, 2005). Higher values reflect more within-network connections. Given that we assigned each thalamic voxel to a cortical functional network in the functional parcellation atlas, higher WMD values reflect more within-network connections of the thalamic voxel with the network it was assigned to (see Methods). We found that thalamic voxels in both first order and higher order thalamic nuclei exhibited higher WMD values than most cortical ROIs (Figure 3A). A two-sample Kolmogorov-Smirnov test showed that on average thalamic voxels had significantly higher WMD values when compared to cortical ROIs (mean WMD for thalamic voxels = 1.62, SD = 1.9; mean WMD for cortical ROIs = 0, SD = −.89; *D_333,2227_* = 0.45, *p* < 10^−10^). We further compared thalamic voxels' WMD values to cortical provincial hubs, which was arbitrary defined as cortical ROIs with WMD values greater than 90% of all cortical ROIs (threshold: WMD = 1.04). We found that on average thalamic voxels exhibited WMD values that were comparable to cortical provincial hubs (mean WMD for cortical provincial hubs = 1.35, SD = 0.26; mean WMD for voxels within first order thalamic nuclei = 1.37, SD = 1.79, mean WMD for voxels within higher order thalamic nuclei mean WMD = 1.75, SD = 1.94). In addition to the voxel-wise analysis, we also calculated WMD values for each thalamic subdivision derived from the functional parcellation atlas (Figure 2A). We found that thalamic subdivisions associated with the cingulo opercular (CO), default mode (DM), frontoparietal (FP), medial temporal (mT), and superior frontoparietal (sFP) networks exhibited higher WMD values when compared to cortical provincial hubs (Figure 3B).

**Figure 3.**
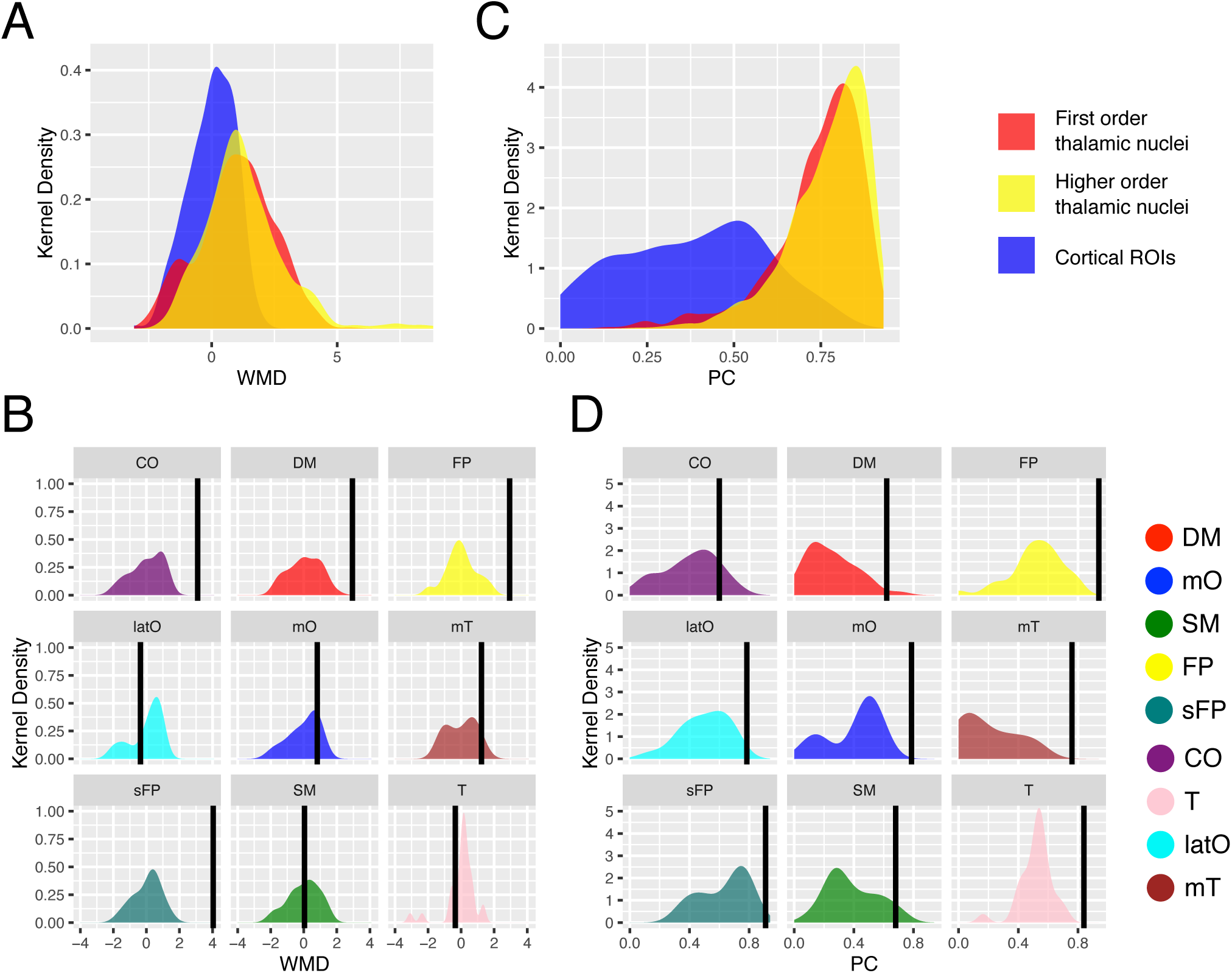
Nodal properties of the thalamus and cortical ROIs. (A) Kernel density plot of WMD values for thalamic voxels and cortical ROIs. Thalamic voxels were categorized into two categories of thalamic nuclei (Sherman, 2016; Sherman and Guillery, 2013). (B) Kernel density plot of cortical WMD values summarized by cortical functional networks. WMD values for each thalamic functional subdivision is represented by the black vertical bar. (C) Kernel density plot of PC values for thalamic voxels and cortical ROIs. Thalamic voxels were categorized into two categories of thalamic nuclei. (D) Kernel density plot of PC values summarized by cortical functional networks. PC values for each thalamic functional subdivision is represented by the 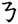 black vertical bar. First order thalamic nuclei included AN, LGN, MGN, VL, and VP. Higher order thalamic nuclei included IL, MD, LP, Po, Pulvinar, VA, and VM. Box plot percentiles. All graph metrics averaged across network densities (see Methods).

#### Connector Hub Property Analyses

Connector hub properties can be measured by the graph theory metric called participation coefficient (PC), which is a measure of the strength of inter-network connectivity for each region normalized by their expected value (Guimera and Amaral, 2005). Higher values reflect more inter-network connections. For each thalamic voxel, we calculated its PC value based on it inter-network connectivity pattern to all cortical ROIs. We found that thalamic voxels in both first order and higher order thalamic nuclei exhibited higher PC values than most cortical ROIs (Figure 3C). A two-sample Kolmogorov-Smirnov test showed that thalamic voxels had significantly higher PC values compared to cortical ROIs (mean PC for thalamic voxels = 0.76, SD = 0.12; mean PC for cortical ROIs = 0.36, SD = 0.22; *D*_*333,2227*_ = 0.65, *p* < 10^−10^). We further compared thalamic voxels' PC values to cortical connector hubs, which defined as cortical ROIs with PC values greater than 90% of all cortical ROIs (threshold: PC = 0.64). We found that on average voxels in both first order and higher order thalamic nuclei exhibited high PC values that were comparable to cortical connector hubs (mean PC for cortical connector hubs = 0.72, SD = 0.05; mean PC for voxels in first order thalamic nuclei = 0.74, SD = 0.13, mean PC for voxels in higher order thalamic nuclei = 0.77, SD = 0.11). We repeated our analysis by calculating PC values for each thalamic functional subdivision derived from the functional parcellation atlas (Figure 2A). We found that all functional thalamic subdivisions exhibited high PC values when compared to cortical ROIs (Figure 3D).

#### Spatial Distribution of Connector and Provincial Hub Properties

We found that cortical ROIs in the precuneus, medial frontal, inferior parietal, insular and middle frontal cortices exhibited high WMD values, whereas ROIs in anterior and inferior frontal, superior precentral sulcus, intraparietal sulcus, and lateral occipital cortices exhibited high PC values (Figure 4A). High PC and WMD values were found throughout the thalamus (Figure 4B-C). To determine differences in the spatial distribution of connector and provincial hub properties in the thalamus, we identified thalamic voxels that exhibited WMD or PC values greater than cortical connector and provincial hubs. We found that anterior, medial, posterior, and dorsal parts of the thalamus exhibited both strong provincial and connector hub properties, whereas portions of the lateral thalamus also exhibited strong connector hub property (Figure 4D). A small portion of the ventral thalamus has only strong provincial hub property.

**Figure 4.**
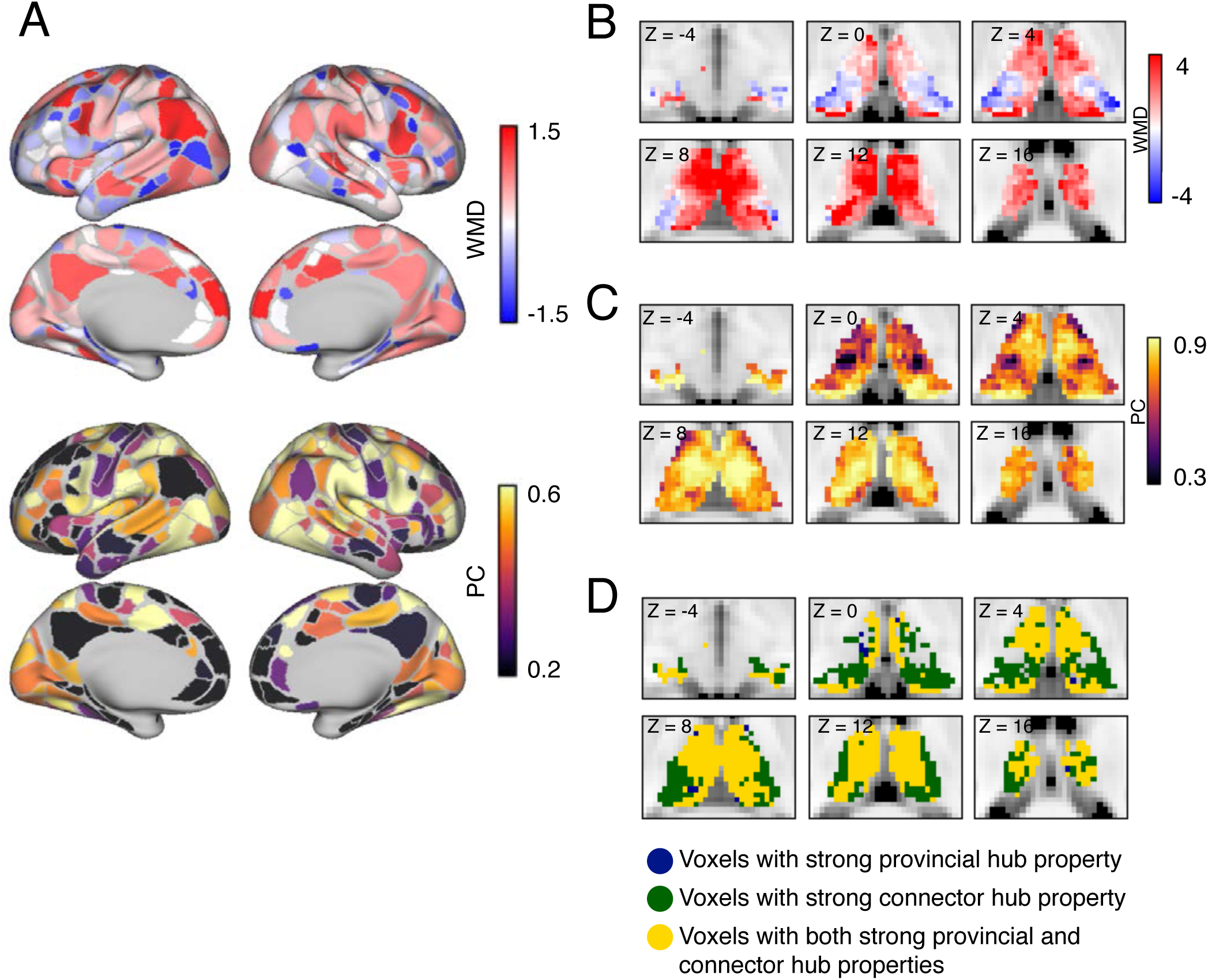
Spatial distribution of network metrics. (A) WMD and PC values of cortical ROIs. (B) WMD values of thalamic voxels. (C) PC values of thalamic voxels. (D) Location of voxels with strong connector (colored in dark navy blue), provincial (colored in dark green), or connector plus provincial hub properties (colored in gold) in the thalamus. In D, Only thalamic voxels that exhibited PC and/or WMD values greater than 90% of all cortical ROIs are displayed. All graph metrics averaged across network densities (see Methods).

#### Connector and Provincial Hub Properties of Each Thalamic Subdivision

We calculated the median WMD and PC values across voxels for each thalamic subdivision, and compared those values to cortical connector and provincial hubs. Based on the functional parcellation atlas, thalamic subdivisions that showed dominant functional coupling with CO, DM, FP, mT, and sFP networks exhibited high WMD values numerically comparable to cortical provincial hubs (Fig 5A). Based on the Oxford-FSL thalamocortical structural connectivity atlas, thalamic subdivisions with dominant structural connectivity with the prefrontal cortex and temporal cortices showed high WMD values comparable to cortical provincial hubs (Figure 5B). Based on the Morel histology atlas, thalamic subdivisions with high WMD values comparable to cortical provincial hubs included the anterior nucleus (AN), LGN, VL, intralaminar nuclei (IL), lateral posterior nucleus (LP), MD, medial pulvinar (PuM), and ventral anterior nucleus (VA) nucleus (Figure 5C). For connector hub properties, we found that all thalamic subdivisions exhibited high PC values comparable or higher than cortical connector hubs (Figure 5D-F). In addition to the voxel-wise analysis, we also calculated graph metrics for each thalamic subdivision. Subdivision-level graph metrics were similar to results from voxel-wise graph metrics (Figure 5A, 5D-F, as indicated by the colored horizontal bar in each individual box-whisker). Note that WMD values were only calculated for the functional parcellation atlas because WMD was defined in relation to a specific cortical functional network, a relation that only exists in the functional parcellation atlas.

**Figure 5.**
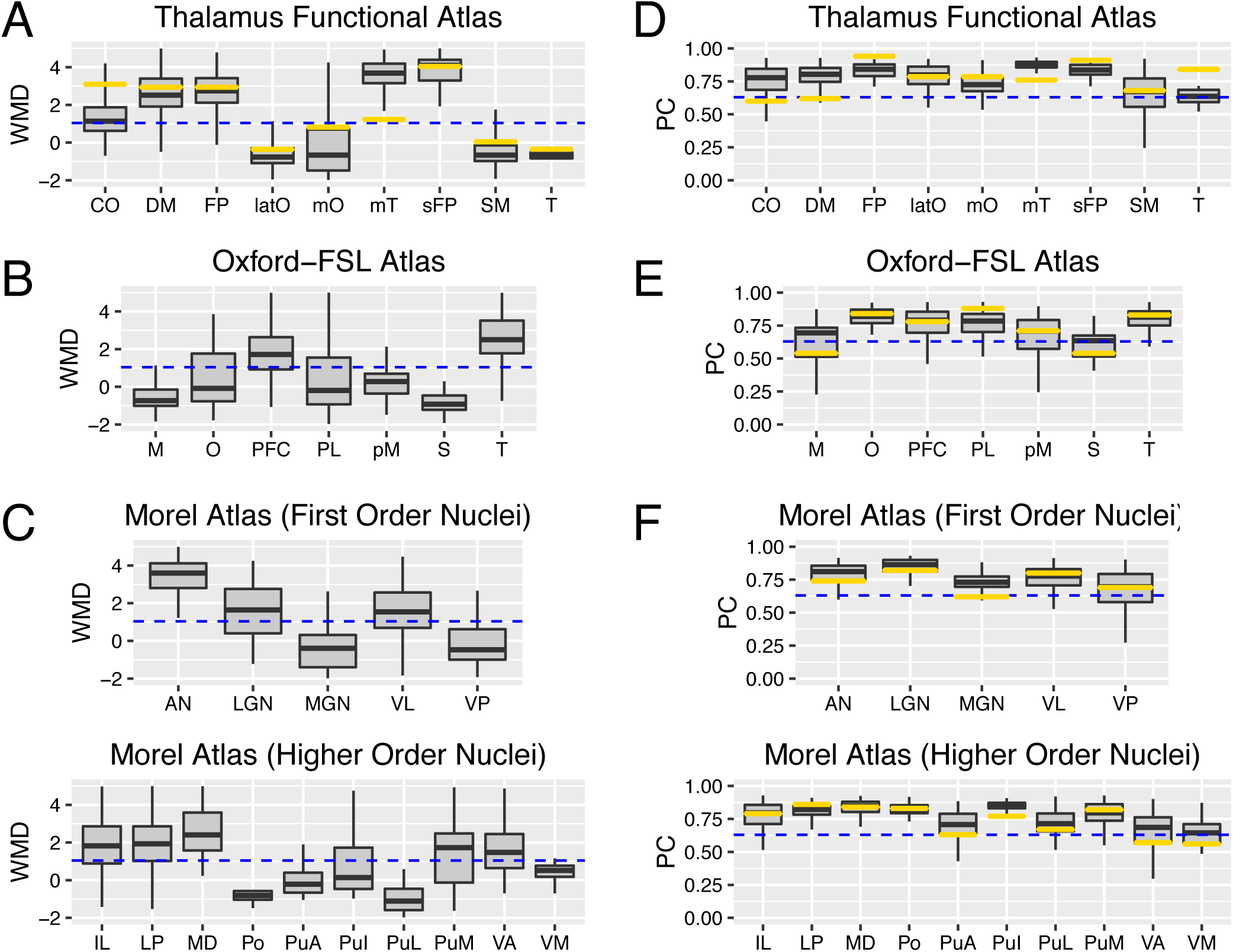
Graph metrics of each thalamic subdivision. (A-C) Box plots summarizing WMD values (A-C) and PC values (D-F) for all thalamic voxels for each thalamic subdivision in each thalamic atlas. The horizontal blue dashed line represents WMD or PC values of cortical provincial or connector hubs (defined as greater than 90% of all cortical ROIs). The horizontal gold bar in each individual box plot represents the graph metrics calculated on the level of thalamic subdivision. Abbreviations for the Oxford-FSL atlas: motor (M), occipital (O), prefrontal (PFC), parietal (PL), premotor (pM), somatosensory (S), temporal (T). Box plot percentiles (5^th^ and 95^th^ for outer whiskers, 25^th^ and 75^th^ for box edges) calculated across voxels for each thalamic subdivision. All graph metrics averaged across network densities (see Methods).

#### Replication of results

We replicated the WMD and PC analyses using an independent rs-fMRI dataset; the spatial correlation values across both cortical ROIs and thalamic voxels between the test and replication datasets for PC and WMD scores were 0.74 (degrees of freedom = 2558, *p* < 10^−5^) and 0.78 (degrees of freedom = 2558, *p* < 10^−5^), respectively. We also replicated our results using a different cortical ROI definition template that consists of 320 cortical ROIs (Craddock et al., 2012),the thalamic voxel-wise spatial correlation values for PC and WMD scores were 0.63 (degrees of freedom = 2225, *p* < 10^−5^) and 0.78 (degrees of freedom = 2225, *p* < 10^−5^), respectively. Finally, to ensure our results were not specific to calculating graph metrics on weighted networks, we recalculated PC values using binary networks, and obtained similar results. The spatial correlation of PC values between weighted and binary networks was 0.98 (degrees of freedom = 2225, *p* < 10^−5^).

### Connectivity Patterns of Specific Thalamic Nuclei

Based on the Morel histology atlas, we found that AN, LGN, VL, IL, LP, MD, PuM, and VA exhibited both strong provincial and connector hub properties comparable to cortical hubs. To further probe their connectivity patterns, for each nucleus we calculated its mean functional connectivity strength with each of the 9 cortical functional networks (using partial correlations, see Methods), and divided by the nucleus's summated total connectivity strength with all networks. If a nucleus is diffusely interacting with all functional networks, then it should devote ∼11% (1/9 = 0.11) of its total connectivity for each network. In contrast, if a nucleus only interacts with a selective network, the majority of its connectivity strength should be devoted to that network, while connectivity with other networks should be considerably lower. Consistent with its strong connector hub properties, we found that each of these thalamic nuclei exhibited a diffuse functional connectivity pattern, with strong connectivity (>11% of its total connectivity strength) with multiple cortical functional networks (Figure 6).

**Figure 6.**
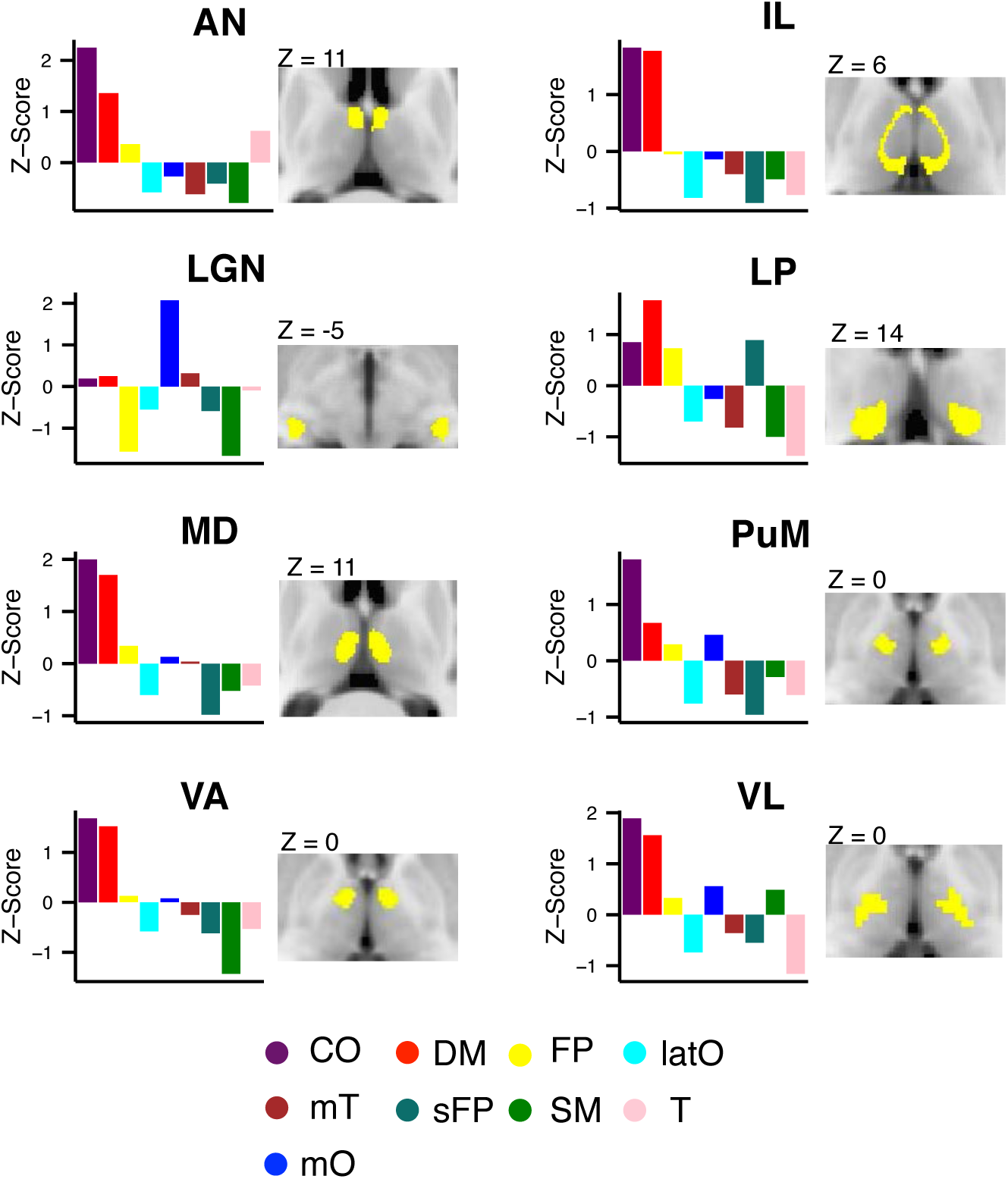
Distributive pattern of thalamocortical connectivity for thalamic nuclei. Cortical functional networks most strongly connected with the following thalamic nuclei: AN, LGN, VL, VA, VM, IL, LP, MD, PuM. Thalamic nuclei (labeled in yellow) are displayed on axial MRI images. The bar graphs represent the distribution of connectivity strength between thalamic nuclei and each of the 9 cortical functional networks. The dashed line represents the expected proportion of total connectivity if connections were equally distributed across networks.

### Meta-Analysis of the BrainMap Database

We found that multiple thalamic subdivisions exhibited both strong provincial and connector hub properties, suggesting that the thalamus is capable of mediating information communication both within and between multiple functional brain networks. Given that individual cortical functional networks are putatively associated with distinct cognitive functions (Bertolero et al., 2015; Smith et al., 2009), it is likely that any individual thalamic nucleus with distributive connectivity with multiple cortical functional networks is involved in multiple cognitive functions. We tested this hypothesis that by analyzing results from a published meta-analysis of 10,449 functional neuroimaging studies (Yeo et al., 2015). This published meta-analysis derived latent variables—an ontology of cognitive functions or “cognitive components”— that best described the relationship between 83 behavioral tasks and corresponding brain activity. From this data, a “cognitive flexibility score” can be estimated by summing the number of cognitive components that are engaged by every brain region (Yeo et al., 2015). Here, a brain region with high cognitive flexibility score is assumed to be involved in more cognitive functions. We have previously demonstrated that cortical connector hubs exhibit high cognitive flexibility scores (Bertolero et al., 2015; Yeo et al., 2016). We hypothesize that thalamic subdivisions that are recruited by multiple cognitive components likely serve as integrative connector hubs, whereas thalamic subdivisions that are recruited by a limited number of specific functions are likely involved in domain general processes.

Consistent with our hypothesis, the thalamus was found to be involved in multiple cognitive components (Figure 7A). A two-sample Kolmogorov-Smirnov test showed that thalamic voxels have higher cognitive flexibility scores when compared to cortical ROIs (mean for thalamic voxels = 3.67, SD = 1.71; mean for cortical ROIs = 0.36, SD = 0.22; *D*_*333,2227*_ = 0.54, *p* < 10^−10^). We further examined the specific cognitive components (C1-C12, see ref 19 for details) that recruited each of the thalamic subdivisions. As an example, VL, with projections to motor and premotor cortices (Alexander et al., 1986), is recruited by components C1 and C2 that predominately recruit motor cortices (Figure 7B). However, VL also participates in other cognitive components that recruit lateral prefrontal, medial prefrontal, and parietal cortices (C8, C9, C12, Figure 7B).

**Figure 7.**
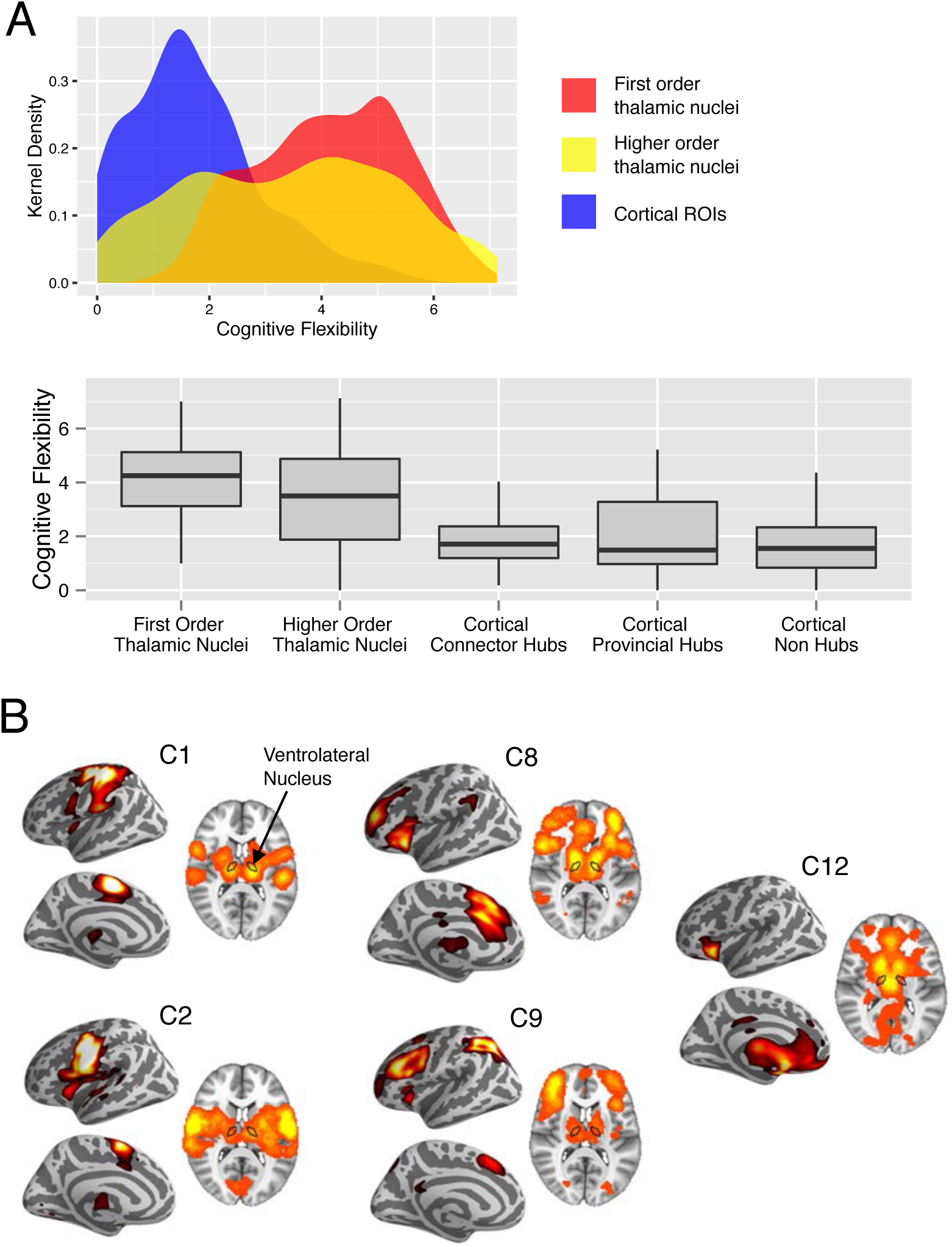
Cognitive flexibility score and cognitive components. (A) kernel density plot and box-plots of cognitive flexibility scores for thalamic voxels and cortical ROIs. Thalamic voxels were categorized into two categories of thalamic nuclei. Box plot percentiles (5^th^ and 95^th^ for outer whiskers, 25^th^ and 75^th^ for box edges) calculated across voxels for each thalamic nuclei type or across cortical ROIs for each cortical ROI type. (B) Spatial distribution of brain activity engaged by each cognitive components recruited by the thalamic nucleus VL.

### Thalamic lesions have Global and Distal Effects on Cortical Network Organization

Modularity is a metric that quantifies the extent to which the brain is differentiated into separable sub-networks, and is an essential property found in many complex systems (Sporns and Betzel, 2016). Based on our findings from a study of patients with focal cortical lesions (Gratton et al., 2012), we predict that if thalamic subdivisions serve as connector hubs for functional brain networks, lesions to those subdivisions should reduce modular organization of these networks. Modularity can be measured by Newman's modularity Q (Newman, 2006), a comparison between the number of connections within a module to the number of connections between modules. In four patients with focal thalamic lesions (Figure 8A; one patient has bilateral lesions, three have unilateral lesions), we examined the effect of a thalamic lesion on cortical modularity across the whole cerebral cortex, the lesioned hemisphere, and the intact hemisphere (Figure 8B). Each patient's Q score was converted to a z-score using the mean and standard deviation of healthy controls (see Methods). In all four patients, whole-brain modularity was lower (as indicated by negative z-scores). In patients with unilateral thalamic lesions, lower modularity was found in both the lesioned and nonlesioned hemispheres, suggesting that a unilateral thalamic lesion impacts modular organization of both hemispheres.

**Figure 8.**
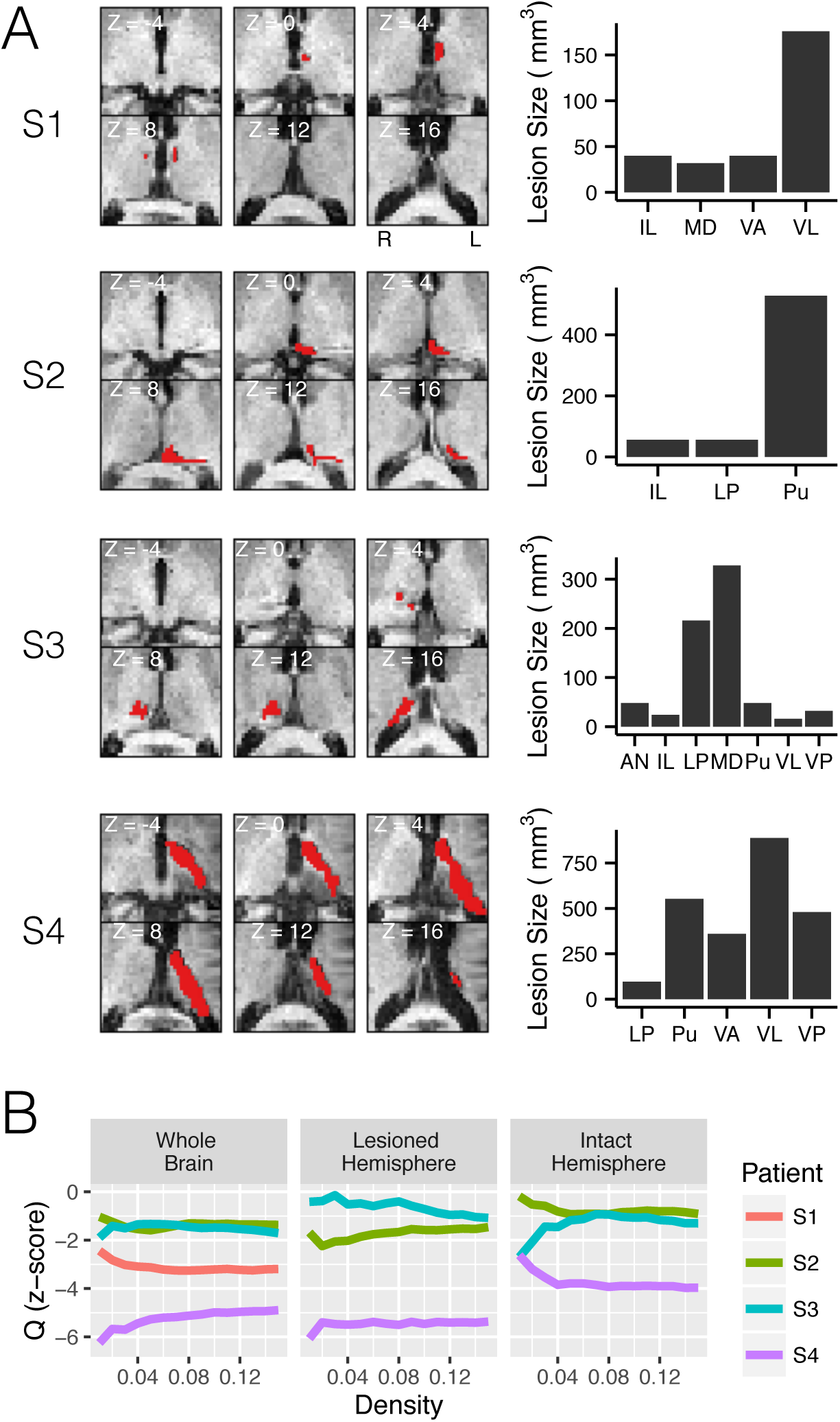
Focal thalamic lesion's effect on cortical network organization. (A). MRI scans of thalamic lesions (marked in red) in four patients (S1-S4). Lesion size for each patient in each thalamic subdivision is summarized in bar graphs. (B) Individual patient's normalized modularity score across network densities, presented separately for the whole cerebral cortex, the lesioned and the nonlesioned hemispheres.

### Discussion

In this study, we provide evidence suggesting that the human thalamus is a critical integrative hub for functional brain networks. First, we found that the thalamus exhibited distributive functional connectivity with multiple cortical regions. Importantly, all thalamic subdivisions exhibited strong between-network connectivity, indicating that a single thalamic subdivision not only connects with multiple cortical regions, but also with multiple cortical functional networks. From a meta-analysis of 10,449 neuroimaging experiments, we further found that the thalamus is engaged by multiple cognitive functions, supporting a role in multimodal information processing. Finally, we found that focal thalamic lesions cause a disruption of the modular structure of cortical functional networks, further underscoring the critical contribution of thalamic function to brain network organization.

The human brain is composed of modular functional networks (Bertolero et al., 2015), which comprise provincial hubs — brain regions important for within network communication — and connector hubs — brain regions important for communication between networks. Here, we used graph-theoretic measures to estimate provincial and connector hub properties of the thalamus. Consistent with traditional interpretations of thalamic function, multiple thalamic subdivisions exhibited strong provincial hub properties. Moreover, both first order and higher order thalamic nuclei exhibited distributive between-network connectivity patterns and strong connector hub properties (Figure 3–6). These results suggest that individual thalamic nuclei, which traditionally were thought to be only connected to a single functional network, in fact have extensive connectivity with multiple functional networks, a pattern that is similar to cortical connector hubs. Cortical connector hubs are proposed to integrate information across segregated functional brain networks (Bertolero et al., 2015; Cole et al., 2013; van den Heuvel and Sporns, 2013; Yeo et al., 2015). Thalamic nuclei's widespread connectivity pattern allows the thalamus to send and access information across diverse cortical functional networks. Via convergence of information, the thalamus may serve as an integrative hub that subserves multiple cognitive functions. Although it has previously been proposed that the thalamus is not simply a relay station but serves to mediate cortical to cortical communication (Sherman, 2016), the notion that the thalamus also plays integrative role interacting with multiple functional brain networks has received less attention in studies of thalamic function.

Higher order thalamic nuclei, which receive inputs predominately from the cortex, are hypothesized to provide trans-thalamic routes to support cortico-cortical interactions within a functional network that receive its projections (Saalmann et al., 2012; Sherman, 2016). For example, the posterior nucleus transfers information from primary area to secondary somatosensory areas (Theyel et al., 2010). Likewise, the pulvinar has extensive reciprocal connections with striate and extrastriate visual cortices (Adams et al., 2000), and is thought to modulate information communication between visual areas (Saalmann and Kastner, 2011; Saalmann et al., 2012). In contrast, first order thalamic nuclei, which receive projections from peripheral sensory organs or other subcortical structures, have projections to primary cortices, and are thought to act as modality-selective relays to relay a limited type of afferent signal to the cortex. Our graph-theoretic analyses of thalamocortical functional connectivity provide evidence suggesting that both first order and higher order thalamic nuclei not only participate in information exchange between cortical regions that they project to, but also further interact with multiple cortical functional networks.

Thalamic nuclei project to and receive projections from multiple brain regions that belong to different functional brain networks. For example, higher order nuclei have higher concentration of “matrix” thalamocortical cells that show diffuse thalamocortical projections unconstrained by the boundaries of cortical topographic representations, and receive non-reciprocal cortico-thalamic innervations from multiple cortical regions (Jones, 2001; McFarland and Haber, 2002). The inhibitory thalamic reticular nucleus, rich in GABAergic cells, also receives input from the cortex and the basal ganglia, and further modulate activity in both first order and higher order thalamic nuclei (Sherman and Guillery, 2013). Thalamic nuclei could also have dense reciprocal connections with cortical connector hubs that in turn are connected with multiple cortical functional networks. A rodent fMRI study further demonstrated that the mouse thalamus has extensive inter-network connectivity with multiple functional sub-networks in the rodent brain (Liska et al., 2015). These findings suggest that thalamocortical functional connectivity has an anatomical substrate capable of simultaneously receiving and transmitting signals between multiple cortical functional networks.

Consistent with its extensive connectivity with multiple cortical functional networks, we found that the thalamus is one of the most “cognitively flexible” brain regions, indicating that the thalamus is involved in a diverse range of behavioral tasks. This observation derived from our meta-analysis of the BrainMap database is further supported by several representative empirical studies demonstrating that the thalamus mediates interactions between higher order cognitive processes (e.g., attention and working memory) and more elementary sensorimotor functions (de Bourbon-Teles et al., 2014; Saalmann et al., 2012; Zhou et al., 2016). For example, a non-human primate electrophysiology study found that deactivating the pulvinar reduced the attentional effects on sensory-driven evoked responses recorded in V4 (Zhou et al., 2016). Also, optogenetically perturbing thalamic activity in rodents impaired animals' ability to select between conflicting visual and auditory stimuli (Wimmer et al., 2015). Finally, VL lesions in humans impair their ability to utilize a memorized cue in working memory to guide visual search of multiple visual stimuli (de Bourbon-Teles et al.,2014). Together, results from our graph analyses of thalamocortical functional connectivity and meta-analysis of task-related thalamic activity patterns suggest that the thalamus participates in interactions between multiple functional cortical networks, networks that are putatively involved in distinct cognitive functions. Based on our empirical results, future task-based studies can test this hypothesis regarding the role of the thalamus in integrative functions.

Previous studies suggest that connector hubs are critical for maintaining the modular architecture of functional brain networks (Gratton et al., 2012). Mathematically, whole brain modularity is inversely related to between-network connectivity, therefore a loss of connector hub should increase modularity. However, consistent with our previous work with patients with focal cortical lesions (Gratton et al., 2012), we found that thalamic lesions reduce cortical functional networks' modularity. This suggests that thalamic connector hub damage not only decreases between-network connectivity, but also alters the large-scale organization of cortical networks. This finding is also consistent with a TMS study in healthy subject that found that disruption of connector hub function increased between-network connectivity (Gratton et al.,2013). Thus, the effect of a thalamic lesion is not constrained only to cortical regions it directly projects to, but further causes “connectomal diaschisis” in cortical functional networks (Carrera and Tononi, 2014). In summary, in addition to playing a role in functional integration of cortical networks, the thalamus is also likely necessary for maintaining the modular structure of functional brain networks.

## Methods

### Datasets

For the main analyses, we analyzed publically available resting-state fMRI (rs-fMRI) data from 303 subjects (mean age = 21.7, STD = 2.87, age range =19-27, 131 males) that were acquired as part of the Brain Genomics Superstruct dataset (Holmes et al., 2015). For each subject, two runs (6.2 minutes each) of rs-fMRI data were collected using a gradient-echo echo-planar imaging sequence with the following parameters: relaxation time (TR) = 3000 ms, echo time (TE) = 30 ms, flip angle = 85 degrees, 3 mm^3^ isotropic voxels with 47 axial slices. Structural data were acquired using a multi-echo T1-weighted magnetization-prepared gradient-echo (MPRAGE) sequence (TR = 2,200 ms, TE= 1.54 ms for image 1 to 7.01 ms for image 4, flip angle = 7 degree, 1.2 mm^3^ isotropic voxel). We replicated our main analyses with publically available rs-fMRI data from 62 healthy adults (mean age = 22.59, SD = 2.45, age range =19-27, 26 males) that were acquired as part of the NKI-Rockland sample (Nooner et al., 2012). For each subject, 9 minutes and 35 seconds of rs-fMRI data were acquired using a multiband gradient-echo echo-planar imaging sequence (TR = 1400 ms, echo time = 30 ms, multiband factor = 4, flip angle = 65 degrees, 2 mm^3^ isotropic voxels with 64 axial slices). Structural data were acquired using a MPRAGE (TR = 1900 ms, TE= 2.51 ms, flip angle = 9 degree, 1 mm^3^ isotropic voxel). For both datasets, subjects were instructed to stay awake and keep their eyes open.

For the lesion analyses, we analyzed rs-fMRI data from four patients with focal thalamic lesions (ages: S1 = 81 years, S2 = 49 years, S3 = 55 years, S4 = 83 years all males, all were scanned at least 6 months after their stroke). Two runs of rs-fMRI data were collected (10 minutes each; TR = 2000 ms, echo time = 30 ms, flip angle = 72 degrees, 3.5 mm^2^ in plane resolution with 34 axial 4.2 mm slices). Structural images were acquired using a MPRAGE sequence (TR = 2,300 msec, TE = 2.98 msec, flip angle = 9°, 1 mm^3^ voxels). Patients were instructed to stay awake and keep their eyes open. Informed consent was obtained from all patients in accordance with procedures approved by the Committees for Protection of Human Subjects at the University of California, Berkeley.

### Functional MRI Data Preprocessing

Image preprocessing was performed the software Configurable Pipeline for the Analysis of Connectomics (Sikka et al., 2014). First brain images were segmented into white matter (WM), gray matter, and cerebral spinal fluid (CSF). Rigid body motion correction was then performed to align each volume to a temporally averaged volume, and a boundary-based registration algorithm was used to register the EPI volumes to the anatomical image. Advanced Normalization Tools (ANTS) was used to register the images to MNI152 template using a nonlinear normalization procedure (Avants et al., 2008). We then performed nuisance regression to further reduce non-neural noise and artifacts. To reduce motion-related artifacts, we used the Friston-24 regressors model during nuisance regression (Friston et al., 1996). WM and CSF signals were regressed using the CompCor approach with five components (Behzadi et al., 2007). Linear and quadratic drifts were also removed. Because the physical proximity between the thalamus and the ventricles could result in blurring of fMRI signal, we further regressed out the mean signal from CSF, WM, and gray matter that were within 5 voxels (10 mm) from the thalamus. After regression, data were bandpass filtered from 0.009–0.08 Hz, and scaled to a whole-brain mode value of 1000. No spatial smoothing was performed.

### Identifying Cortical Functional Networks

Following preprocessing, mean rs-fMRI time-series were extracted from 333 cortical ROIs (Gordon et al., 2016), and concatenated across runs for subjects with multiple rs-fMRI scans. Cortico-cortical functional connectivity was assessed in each subject by computing Pearson correlations between all pairs of cortical ROIs, resulting in a 333 x 333 correlation matrix. This correlation matrix was then thresholded at different density thresholds to retain the strongest percentages of functional connections. For each subject, putative cortical functional networks were then identified in individuals and then average across subjects using a recursive InfoMap algorithm to partition the matrix into modules by integrating results across thresholds (Bertolero et al., 2015). We started by identifying networks in individuals. First, we identified functional networks using the InfoMap algorithm (Rosvall and Bergstrom, 2007) at a threshold of density =0.15. A consensus matrix (a value of 1 where the two ROIs are in the same module and a value of 0 elsewhere; (Lancichinetti and Fortunato, 2012) was formed based on this result. The threshold was then decreased by density = 0.001, and the InfoMap community detection was ran again. The consensus matrix was then updated for the new partition, except for rows and columns for which the node had no edges in the current version of the graph or the node was not in a network with at least 5 ROIs. This procedure continued until to the minimal threshold of density = 0.01. Thus, the consensus matrix represents the community assignments for each pair of nodes at their sparsest level possible (i.e., before they become disconnected from the graph). We then aggregated individual subjects' module organization by averaging these consensus matrices across subjects. This averaged matrix was then submitted to the same recursive method to identify group-level cortical functional networks. Networks with 5 or fewer ROIs were eliminated from further analyses, and ROIs that were not clustered into networks were excluded from further graph analyses.

### Thalamus Parcellation

To localize the thalamus, the Morel Atlas (Krauth et al., 2010) was used to define its spatial extent (2227 2 mm^3^ voxels included in the atlas). To identify thalamic subdivisions, three different thalamic atlases were utilized. We first performed a custom winner take all functional parcellation using rs-fMRI data. We calculated partial correlations between the mean BOLD signal of each cortical functional network and the signal in each thalamic voxel, while partialing out signal variance from other functional networks. Partial correlations were then averaged across subjects, and each thalamic voxel was labeled according to the cortical network with the highest partial correlation. The Morel atlas identified thalamic nuclei based on cyto- and myelo-architecture in stained slices of post-mortem tissue collected from five postmortem brains (Morel et al., 1997), and further transformed to MNI space (Krauth et al.,2010). The Oxford-FSL thalamic structural connectivity atlas defined thalamic subdivisions based on its structural connectivity with different cortical regions estimated from diffusion imaging data (Behrens et al., 2003).

### Thalamic and Cortical Nodal Properties

To formally quantify the network properties of thalamocortical functional connectivity, for each subject, we first extracted signal from the thalamus by either taking each voxel's preprocessed signal, or averaged voxel-wise BOLD signal within each thalamic subdivision. We then calculated the partial correlation between the mean BOLD signal of each cortical ROI and thalamic voxel or subdivision. Partial correlation was calculated by partialing out signal variance from all other cortical ROIs. Given the large number of cortical ROIs, a dimension reduction procedure using principal component analysis was performed on signals from cortical ROIs not included in the partial correlation calculation, and eigenvectors that explained 95% of variance were entered as additional nuisance regressors in the model. We chose partial correlations over full correlations because past studies have shown that detailed thalamocortical connectivity patterns could be obscured without accounting for shared variance between cortical regions (Zhang et al., 2008). Note that no correlations were calculated between thalamic voxels. This resulted in two thalamocortical connectivity matrices: voxel-wise and subdivision-wise. Thalamocortical connectivity matrices were then averaged across subjects, thresholded by network density to retain top 1% to top 15% strongest percentages of functional connections, and submitted to graph analyses. For cortical ROIs, we calculated Pearson correlations between all pair-wise cortical ROIs to obtain cortico-cortical connectivity matrices. Matrices were averaged across subjects and submitted to graph analyses. All graph metrics were calculated across a range of density thresholds.

For each thalamic voxel, subdivision, and cortical ROI, we calculated participation coefficient (PC) and within module degree (WMD) (Guimera and Amaral, 2005). PC value region *i* is defined as:

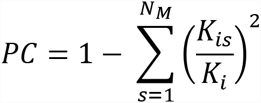

where *k*_*i*_ is the sum of connectivity weight of *i* and *K*_*is*_ is the sum of connectivity weight between *i* and cortical network *s*. N_M_ is the total number of networks. If a region has connections uniformly distributed to all cortical networks, then its PC value will be close to 1; on the other hand, if its connectivity is concentrated within a specific cortical network, its PC value will be close to 0. We further divided PC values by its theoretical upper limit based on the number of functional networks, so that the highest possible PC value given the network architecture would be 1. For comparison, we also calculated PC using binary networks by setting weights above the threshold to 1.

To calculate WMD, correlation matrices were first binarized by setting weights above the threshold to 1. Weights were binarized to equate the connectivity weights between thalamocortical and cortico-cortical networks. WMD is calculated as:

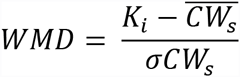

where 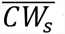 is the average number of connections between all cortical ROIs within cortical network *s*, and *σCW*_*s*_ is the standard deviation of the number of connections of all ROIs in network *s*. *K*_*i*_ is the number of connections between *i* and all cortical ROIs in network *s*. Because our goal was to understand the thalamus's contribution to cortical network organization, thalamus's WMD scores were calculated using the mean and standard deviation of within-network degree (number of intra-network connections) for each cortical functional network.

For all patients, rs-fMRI volumes with framewise displacement (FD) that exceeded 0.5 mm were removed from further analysis (scrubbed) after band-pass filtering (mean percentage of frames scrubbed for patients = 13.34%, SD = 9.32%). Lesion masks were manually traced in the native space according to visible damage on a T1-weighted anatomical scan, and further guided by hyperintensities on a T2-weighted FLAIR image. Lesion masks were then warped into the MNI space using the same non-linear registration parameters calculated during preprocessing.

### Modularity

Modularity can be measured by Newman's modularity Q (Newman, 2006), defined as:

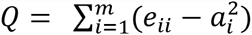

Where *e*_*ii*_ is the fraction of connectivity weight connecting ROIs within a cortical functional network *i*, *a*_*i*_ is the fraction of connectivity weight connecting ROIs in cortical functional network *i* to other cortical networks, and *m* is the total number of cortical functional networks. Network partition from Figure 2A was used. Modularity was calculated for the whole-brain (including all cortical ROIs) and each individual hemisphere by constructing independent networks restricted to only one hemisphere. Note no thalamic subdivision was included into this analysis. For lesions analyses, the mean and standard deviation of Q calculated from all healthy controls were used to normalize Q values (whole brain, damaged and intact hemispheres) from each patient (Gratton et al., 2012):

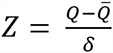

Q is the patient ‘smodurlaity score, 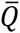 is the average modularity score of healthy controls, and *δ* is the standard deviation in modularity of healthy controls. Modularity calculations were performed separately for each density threshold (from 0.01 to 0.15).

### Meta-Analysis of Functional Neuroimaging Experiments in the BrainMap Database

We reanalyzed data presented in a previously published meta-analysis of the BrainMap database (Yeo et al., 2015). In the meta-analysis, a hierarchical Bayesian model was used to derive a set of 12 cognitive components that best describe the relationship between behavioral tasks and patterns of brain activity in the BrainMap database (Laird et al., 2005). Specifically, each behavioral task (e.g., Stroop, stop-signal task, finger tapping) engages multiple cognitive components, and in turn each cognitive component is supported by a distributed set of brain regions. To determine whether or not a thalamic voxel is recruited by a cognitive component, a threshold of *p* = 10^−5^ was used. This is an arbitrary yet stringent threshold that was used in two prior studies (Bertolero et al., 2015; Yeo et al., 2015). Critically, there is potential spatial overlap between components. Therefore, brain regions that can flexibly participate in multiple cognitive components could be identified by calculating the number of cognitive components each brain region engages. The number of cognitive components was summed for each voxel and cortical ROIs, and defined as a “cognitive flexibility” score (Yeo et al., 2015).

## Acknowledgements

This work was supported by NIH Grants RO1 NS79698 and F32 NS090757. The Morel atlas was obtained by a written consent with Professor G. Szekely from the Computer Vision Laboratory of ETH Zurich, Switzerland. This project was made possible by a collaborative agreement allowing comprehensive access to the BrainMap database, a copyrighted electronic compilation owned by the University of Texas Board of Regents. BrainMap is supported by National Institutes of Health, National Institute of Mental Health Award R01 MH074457.

